# Over half of the phylogenetic diversity accumulated on the African plant tree of life may be eroded under current biodiversity crisis even though threatened species are not evolutionarily unique

**DOI:** 10.1101/2025.02.12.637779

**Authors:** Makuete A.P. Tiawoun, Bopaki Phogole, Bahati Samuel Kandolo, Kowiyou Yessoufou

## Abstract

Although Africa contributes tremendously to global biodiversity, we have a poor understanding of how the African tree of life might be pruned owing to the ongoing biodiversity crisis. Here, we investigated this question integrating statistics and phylogenetics of ∼24000 African vascular plants. We found that 54% of African plant families are hot nodes of threatened species, thus heightening the risk of losing entire clades. We also found that, if all threatened species go extinction, 59% of the evolutionary history of the African plant tree of life would be eroded, and this loss is more than expected at random, although threatened species are not evolutionarily unique. Unfortunately, ∼72% of threatened species and ∼79% of the top-1000-EDGE species are not found in any form of protected areas. Overall, our analysis reveals the extent of biodiversity crisis in Africa and the need for steadfast commitments to i) increased data collection efforts particularly in central African regions and ii) implementation of existing policy for an accelerated recovery.

## 1. Introduction

Globally, over 40,000 species are currently identified as threatened, and nearly 1 million plant and animal species face extinction threats due to anthropogenic pressures [1,2]. The accelerated global biodiversity crisis [3,4] required robust, quick and urgent species extinction risk assessments if we are to bend the curve of biodiversity loss [5] and meet the ambitious goals and targets set to accelerate recovery and ensure a continued provision of human needs [6–8].

Unfortunately, our knowledge of the extent of the crisis is biased toward vertebrates with plants remaining comparatively not only poorly assessed but we have limited knowledge of how the plant tree of life would be pruned under the crisis. For example, we know that 10–30 % of vertebrates, specifically mammals, amphibians and birds are at risk [9], with a third of known amphibians at highest risk among vertebrates [10]. An early report indicated that we have already lost 12% of continental birds and 20% of mammals [11]. In their early report, IUCN [12] indicated that 70% of red-listed angiosperms is at risk of extinction. However, this *a priori* alarming report must be taken with caution because it was based on a small proportion of known plant taxonomic diversity (13000 plants; [12]). Although assessment efforts for plants have increased over the past decade, the efforts remain, however, very limited (62 666 species out of c. 350 000 species, ∼18%; [1]), and controversies around taxonomic, geographic and trait accuracies have been raised against plant risk assessment [13].

Indeed, it is challenging to assess plants due to their ubiquity [14], diversity (∼350 000 plants; [15], data gaps and bias [16]. These challenges prevent a comprehensive risk assessment for all plants, although the Target 2 of the Global Strategy for Plant Conservation made it clear that such comprehensive assessment is a necessity [17]. In the face of the ongoing biodiversity crisis, this knowledge gap must be filled [13,18]. Unfortunately, the challenges are even greater in some regions, e.g., Africa, that are exceptionally rich in biodiversity but characterised by huge knowledge gap [19,20]. Indeed, Africa, representing only 20% of the world’s land, harbours a unique biodiversity. Specifically, Africa hosts the world’s highest extratropical plant diversity including endemic species and theatres of explosive species radiations (e.g. Cape Floristic Region; Madagascar) as well as a relatively intact Pleistocene megaherbivores diversity [20]. Furthermore, Africa hosts about a sixth of global plant richness, ∼17% of mammal richness, 2500 birds, 950 amphibians, about 2000 reptiles and 5000 freshwater fish species [21]. The uniqueness of Africa is also mirrored in the eight (08) global biodiversity hotspots (out of 36) found within its borders [2]. Additionally, the African continent is home to 20% of the world’s rainforest areas, e.g., the Congo Basin, which covers 240-million-hectare of rainforests, supports the livelihoods of 100 million people in the central African region alone and absorbs 4% of global carbon emissions annually [22].

However, nearly one-third of Africa’s vascular plant species is at risk of extinction [23]. Rapid population growth, projected to quadruple by 2100, further exacerbates pressures on African biodiversity [20]. Urban expansion, agricultural intensification, and infrastructure development are leading drivers of habitat fragmentation and loss, intensifying these threats. While protected areas (PAs) play a crucial role in mitigating threats to biodiversity, only 13% of the African continent is designated as PAs [24]. Their effectiveness is often limited by inadequate management and insufficient funding. Even non-protected areas, which face greater pressures from deforestation, agriculture, and illegal activities, also support substantial biodiversity, highlighting the need for conservation strategies that address both protected and non-protected landscapes [25,26].

In the context of global environmental change, identifying the extent of biodiversity loss and determining which species are most at risk of extinction is therefore critical to inform decision-making and prioritizing conservation efforts [27,28]. Traditional approaches to assessing extinction risks, such as field surveys and expert opinions, have provided valuable insights into the vulnerability of African tree species [29]. However, these methods are limited by resource constraints and incomplete data [30]. The International Union for Conservation of Nature (IUCN) Red List (RL) remains the global gold standard for extinction risk assessment. Unfortunately, relying solely on the systematic methodological approach set by IUCN is not producing expected outcomes at a speed that matches that of species loss: only ∼18% of plant species is assessed [1], whilst biodiversity is lost at an exponential rate [31,32]. Only roughly 37 000 African species have been assessed following IUCN methods [33] and 4134 African plant and 1901 animal species are listed under CITES as being threatened with extinction (https://www.speciesplus.net/; accessed 2024).

Interestingly, several attempts have been made to improve the risk assessment following the Red-List systematic approach. These include the Sampled Red List Index [34], the country-level risk assessments [35], the 30/30 challenge which is focused on area-based conservation approach [36,37], modern methodological advances (e.g., criteria-specific and category-predictive approaches [38,39], targeted fundings [40], and availability of custom tools [41,42]. However, these recent attempts to fast-track biodiversity risk assessments have yet to be applied on a large scale (e.g., continental) for large clades such as plants [43], particularly in the developing but species-rich regions of the world where huge biodiversity knowledge gaps are reported (e.g., [19,20]).

This knowledge gap is even widest when it comes to the phylogenetic aspects of the African biodiversity. For example, there is a very scant information about how the current extinction crisis might deplete the evolutionary history or phylogenetic diversity (PD) accumulated over millions of years on the African tree of life. The only study, to our knowledge, that tackles this question on African biodiversity, is one of our studies on African fish family Cyprinidae where we reported that some African lineages are at higher risk than expected and that the loss of all threatened species would lead to a disproportionate loss of PD [44]. On the African flora, such knowledge is required to inform concerted efforts to protect in a comprehensive way the unique evolutionary diversity of the African flora. The need to focus on evolutionary diversity [45–50] is justified by what we now know that the loss of PD would bear more dramatic consequences to biodiversity than that of species richness (see [46,51]). It is further justified by our knowledge that preserving evolutionary diversity is critical to the preservation of a diversity of functions or services [46,52], but also of evolutionarily distinct species [53,54], which may ensure the future provision of feature diversity [52,55,56].

Overall, addressing the complex challenges of biodiversity loss in Africa requires a multi-faceted approach that combines diverse methods for more accurate and comprehensive conservation assessments. One way is to integrate machine learning and phylogenetic analyses to offer a more holistic and innovative framework to address conservation challenges. Machine learning algorithms, particularly neural networks, are effective in predicting extinction risks by analyzing complex datasets, such as species occurrence, climate variables, and human impact data [57,58]. For instance, deep learning techniques have shown promise in predicting species distributions and vulnerability, even in data-deficient regions [59–62]. Phylogenetic analyses enable the identification of evolutionarily distinct species that are vital for ecosystem stability and resilience yet often overlooked in traditional conservation efforts. Studies such as Yessoufou [63] and Tucker [64] highlight the importance of prioritizing these species in conservation planning to preserve both biodiversity and ecosystem services. Cadotte [65] applied phylogenetic analyses to identify key species whose loss might significantly affect ecosystem stability, highlighting the value of focusing on evolutionarily unique species in conservation efforts.

In the present study, our aim is to reveal the extent and consequences of extinction risk in the African flora. First, we employed neural network models to predict extinction risk in the African flora, then identified taxa that are hot spots of threatened species and finally investigated how the loss of at-risk African plant species might impact the evolutionary diversity accumulated over million years on the African plant tree of life.

## 2. Materials and methods

### 2.1. Dataset Compilation

A comprehensive dataset of vascular plant occurrences across the African continent was compiled by retrieving all records (Figure S1) from the Global Biodiversity Information Facility (GBIF) (GBIF.org 2024; accessed October 2024) [66] and the South African National Biodiversity Institute (SANBI) databases (http://redlist.sanbi.org/, accessed October 2024) [67]. This exercise reveals concentrated sampling efforts in southern African regions particularly in South Africa and in species-rich regions such as Mediterranean basin, western Africa forests and eastern Afromontane which are known as African biodiversity hotspots (Figure S1). It also revealed a poor sampling in the Congo basin region which is referred to the African Amazonia (Figure S1).

The dataset that we retrieved also included georeferenced records for all identified taxa from 2020 to 2024. To ensure spatial accuracy, only records with precise geographic coordinates were retained. Since species occurrence records from public databases are often error-prone [27,68,69], the raw data were checked to exclude species with missing geographic coordinates. Following recommendations from prior research for reliable range estimates in conservation assessments, species with fewer than 15 georeferenced records were excluded [27,58,70]. This resulted in a final dataset consisted of 1 162 937 occurrence points, covering 23 362 species. We further retrieved the IUCN status of each of these species from GBIF and SANBI and this was double-checked on the IUCN database. As a result, among the species with at least 15 occurrence points, we identified 6 751 species that are “Data Deficient” (DD; 27 species) and “Non-Evaluated” (NE; 6 724 species). We also identified 1711 species of known risk status, including Least Concern (LC; 1 380 species), Near Threatened (NT; 122), Vulnerable (VU; 101), Endangered (EN; 83), and Critically Endangered (CR; 25). These 1 711 species with at least 15 occurrence points were used to train supervised deep learning models.

### 2.2. Data Analysis

Data analysis was performed combining R (R Development core Team 2021) and Python following Zizka et al.’s [71] approach, with tools such as TensorFlow, Keras, and scikit-learn for machine learning, and the IUCNN R package for feature extraction. Scripts and code necessary to reproduce the analyses are available in public repository (see dataset at: 10.6084/m9.figshare.28390718).

#### Label and Feature Generation

One type of label was generated: 5-Class Prediction, with classes LC, NT, VU, EN, and CR, and these categories were mapped to integer values and transformed into one-hot encoded formats suitable for model training: LC was assigned 0, NT = 1, VU = 2, EN = 3, and CR = 4. Features were derived from species occurrence records using IUCNN R package [72]. These features included geographic variables (e.g., mean latitude and longitude, extent of occurrence, area of occurrence, number of occurrences), climate proxies (e.g., means of 19 bioclimatic variables [73]), and combinations of these features (Tables S1&S2). Species with incomplete feature sets were excluded to ensure compatibility with IUCNN functions. The final dataset comprised 1 711 species with complete features and labels for training, and 6 751 unassessed species for which extinction risk was predicted.

#### Model training, testing, validation and prediction of extinction risk

Neural network classifiers (nn-class) were developed for multiclass classification using TensorFlow, Keras, and scikit-learn [74]. The model architecture consisted of three hidden layers with 71, 46, 5 nodes for the 5-class, each employing Rectified Linear Unit (ReLU) activation functions, and an output layer with 5 neurons, one per class using the Softmax activation function. L2 regularization was applied to prevent overfitting, and the model was compiled with the Adam optimizer [75] and categorical cross-entropy loss. Feature values were normalized using Standard Scaler [74] to ensure equal contribution to the model and split into training (80%) and validation (20%) sets. Synthetic Minority Oversampling Technique (SMOTE) was used to address class imbalance in the training set [76]. Training was conducted for 175 epochs with a dropout rate of 0.1 applied to reduce overfitting and estimate prediction uncertainty [77]. The effective trained models were then used to predict the conservation status of unassessed species (6751 NE and DD species) based on the optimal trained models which were chosen based on their highest validation accuracy.

##### Taxonomic analysis of extinction risk

This analysis was run to determine whether some taxa, e.g., plant families, are particularly targeted by threats. To this end, we first fitted a generalized linear model with a Poisson error distribution to model the number of threatened species in a family as a function of the size of that family. However, in this analysis, we found evidence of data over-dispersion. We therefore developed a negative binomial model to account for over-dispersion. Following this modelling exercise, plant families with positive residuals are considered over-threatened: these are the families that contain more threatened species than would be expected from the fit of the model.

#### Phylogenetic analysis of the extinction risk

##### Assembling the tree of life for African flora

The tree of life of the African flora (23 362 species in our dataset) was reconstructed using the R package ‘V.PhyloMaker’ [78]. First, a list of all the 23 362 species was formed with indication in different columns of the species, genus, and family names as recommended [78]. Then, using the R function ‘phylo.maker’, the phylogenetic tree including all species was reconstructed based on the mega-tree ‘GBOTB.extended.tre’ [79,80]. We referred to the resulting phylogenetic tree of African tree of life.

##### Impacts of species loss on the African tree of Life

We firstly quantified the total phylogenetic diversity PD_total_ on the tree [53] and then calculated PD_threatened_ corresponding to the at-risk PD if all threatened species go extinct. Next, we simulated the loss of all threatened species (12647 species) from the tree by pruning randomly the number of threatened species 1000 times from the tree. For each random pruning, we calculated the corresponding PD lost termed PD_random_loss_. This allowed us to compare the average PD_random_loss_ to PD_threatened_.

##### African plant species on the EDGE: Evolutionary Distinct and Globally Endangered species

We determined the evolutionary distinctiveness (ED; Isaac et al. [81]) for each species using the R function ed.calc (library ape). Using one-way ANOVA, we tested the correlation between ED (after log-transformation) and IUCN categories of species. We repeated the analysis. Finally, we determined the EDGE score for each species (EDGE = [ln(1 + ED) + GE*ln(2)], where GE = global endangerment was coded as follows: LC = 0, NT = 1, VU = 2, EN = 3, CR = 4. We ranked all species based on their EDGE scores and then overlaid the spatial distribution of top 1000-EDGE species over the network of protected areas in Africa.

## 3. RESULTS

### Performance of machine learning models

Our models show validation accuracy of 82.3% and a testing accuracy of 67% (Figure S2a,b). The models predicted LC species with highest precision (0.91) and the other classes with lower precision (Table S3). This is reflected in the confusion matrix (Figure S3) where diagonal elements represent accurate predictions and the off-diagonal elements reveal misclassifications, identifying areas where the model can improve. The conservation status of unassessed species was predicted using the best-trained models, which combined geographic and climate features with three hidden layers and dropout regularization. At the detailed classification level (Figure 1), the model predicted that 52.1% (3582) of the 6717 NE species (Figure 1a) were categorized as Least Concern (LC), followed by smaller proportions classified as Vulnerable (VU), Endangered (EN), Near Threatened (NT), and Critically Endangered (CR). Similarly, for the 27 Data Deficient (DD) species, the model classified the majority (74%, 20 species) as LC, while the remaining 26% were distributed among the VU, EN, NT, and CR categories (Figure 1b).

**Figure 1.**
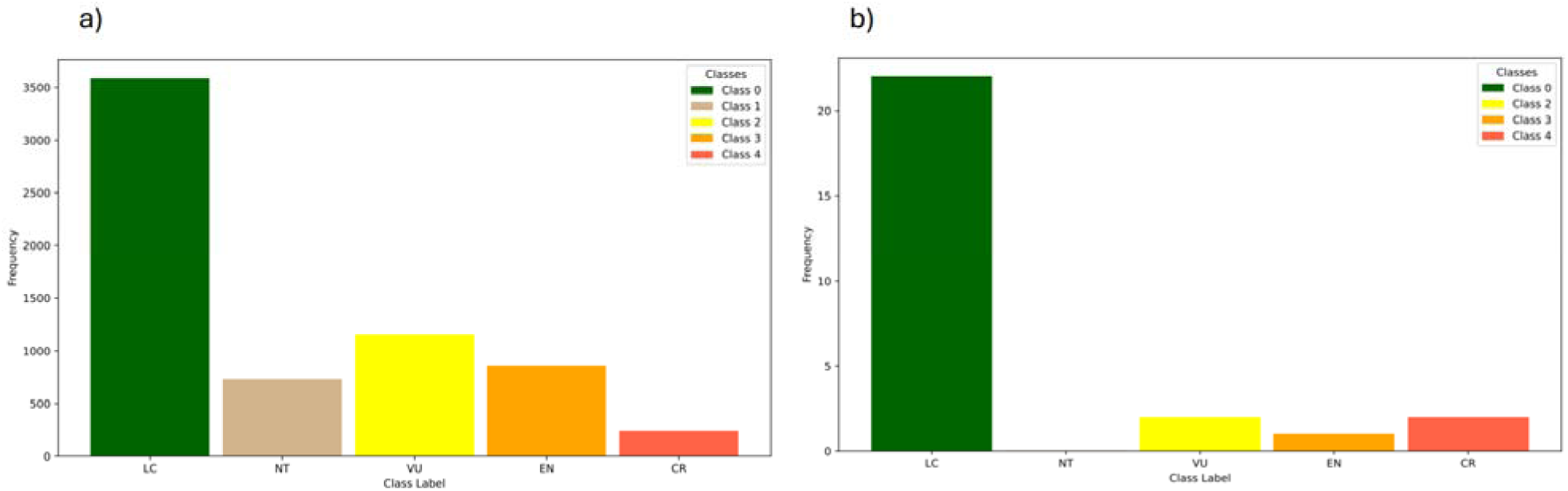
Predicted Detailed Conservation Status of NE and DD Species. (a) Predicted IUCN categories for Not Evaluated (NE) species; (b) Predicted IUCN categories for Data Deficient (DD) species.

### Taxonomic and phylogenetic consequences of extinction risk

We found that 54% of all families (171/316) contain more threatened species than expected (Figure 2; Table S4). The top 4 of such families are Begoniaceae, Bruniaceae, Anacampserotaceae and Zamiaceae (Table S4). This implies that threats to biodiversity target some clades, potentially heightening the risk of losing entire clades. What is the phylogenetic consequence of this? We found that if all threatened species go extinction, this would be equivalent to the loss of 59% of the phylogenetic diversity (PD) accumulated in the African plant tree of life. Also, our simulations of random loss indicate that, if threatened species go extinct, we would lose more PD than expected at random (actual PD loss = 179194.2my vs. mean random expected loss = 139083.2my; CI = [136766.0; 141371.8]; Figure 3).

**Figure 2.**
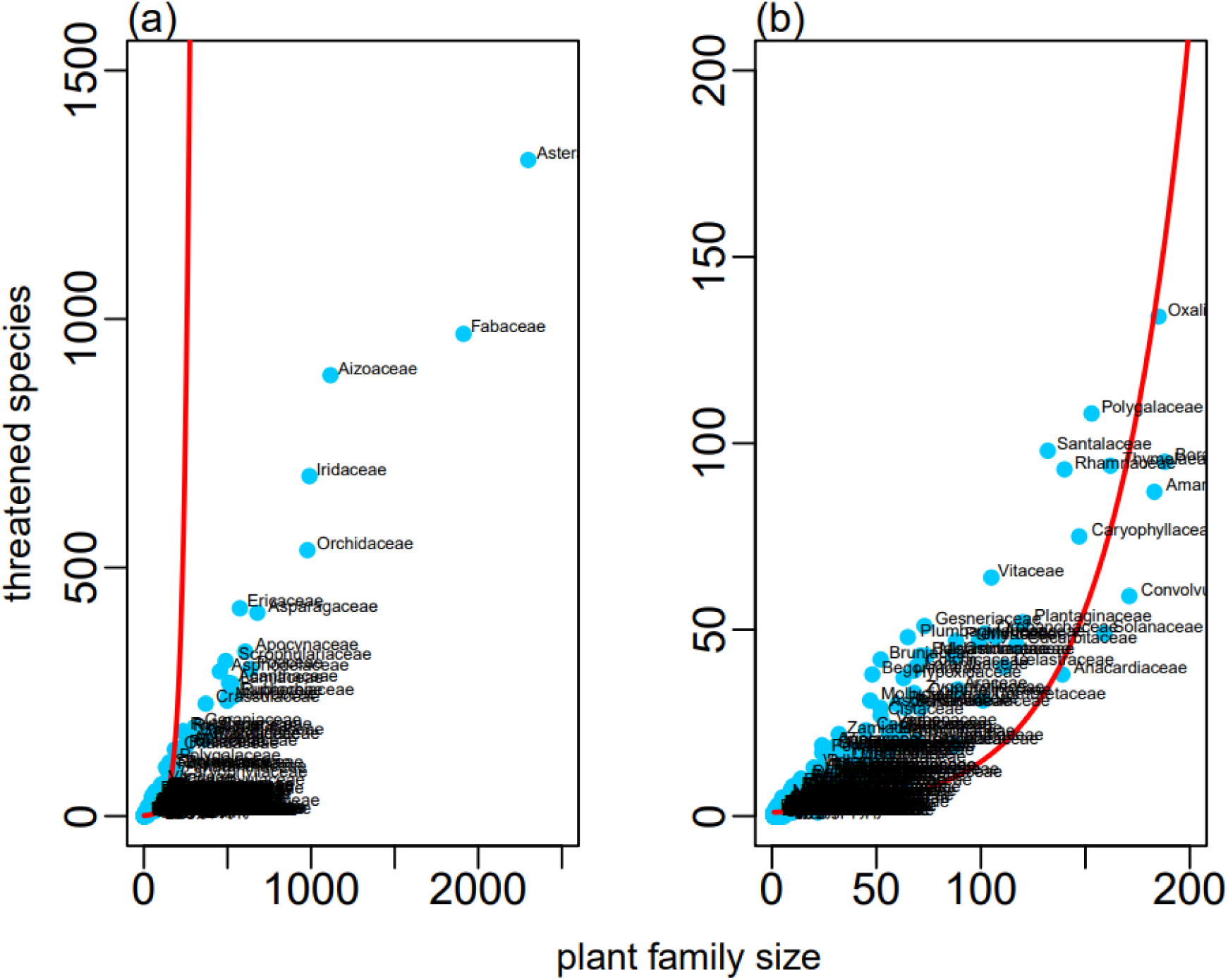
The number of threatened species is dependent on the size of the plant families. (a) negative binomial model illustrating the relationships on the entire dataset; (b) inset showing the details for 200 by 200.

**Figure 3.**
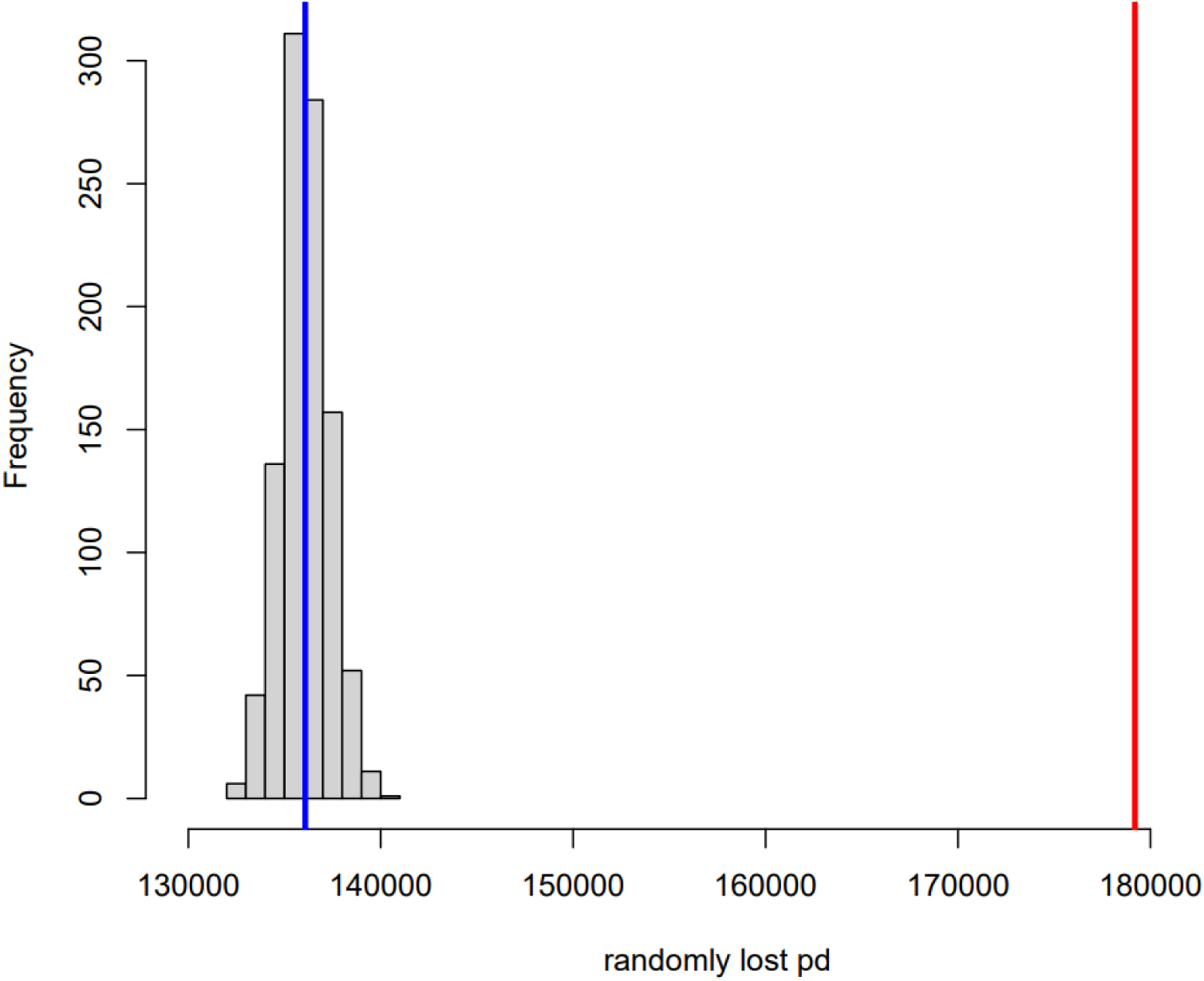
Comparison of observed (red line) and random losses (blue line) of phylogenetic diversity (PD) from the African plant tree of life.

Interestingly, threatened species are not evolutionarily unique species both at 5-threat class (anova; df = 4; P<0.001; Figure 4A) and binary class (anova; df = 1; P<0.001; Figure 4B). Unfortunately, ∼72% of threatened species (Figure S4A) and ∼79% of the top-1000 EDGE species (Figure S4B) are not found in any form of protected areas on the continent. In addition, we ranked species according to their EDGE score (Table S6). *Ptisana robusta* (Marattiaceae), *Rhizoglossum bergianum* (Orchidaceae), *Angiopteris evecta* (Marattiaceae), *A. madagascariensis* (Marattiaceae) and *Philodendron hederaceum* (Oleaceae) are the top 5 most ranked species (in this order) based on EDGE score (Table S5).

**Figure 4.**
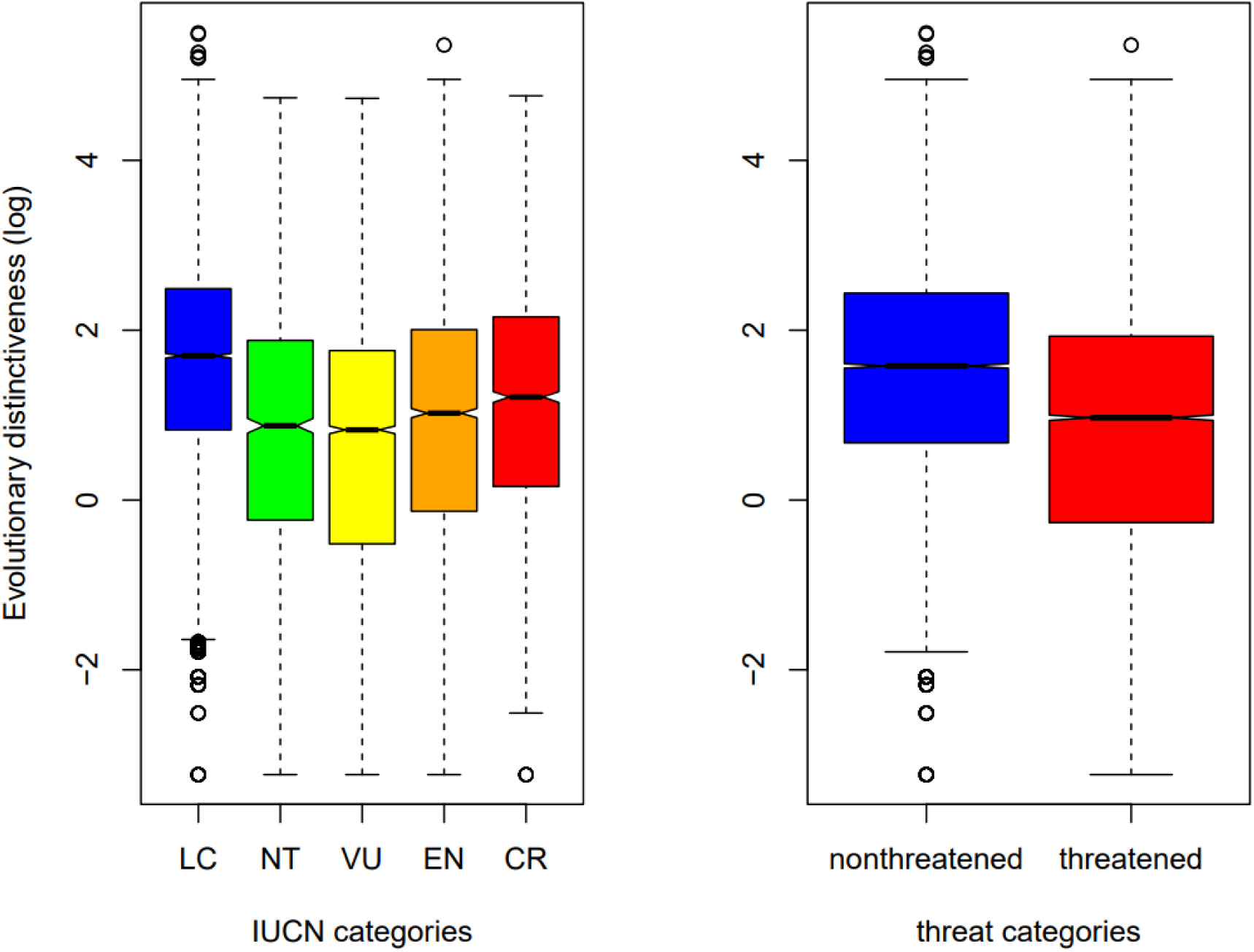
Variation in evolutionary distinctiveness across species of the African plant tree of life for 5-threat classes (A) and binary threat class (B). IUCN categories are Least Concern (LC), Near Threatened (NT), Vulnerable (VU), Endangered (EN), Critically Endangered (CR). Threat categories comprised threatened (VU + EN + CR) and non-threatened categories (LC + NT).

Finally, we identified regions of high-EDGE species. The geography of these regions matches that of African biodiversity hotspots including Guinean forests of West Africa, Mediterranean basin, Eastern Afromontane, Madagascar and succulent Karoo biodiversity hotspots (Figure 5). We additionally identified some countries harbouring high-EDGE species mostly found in the southern African region, including northern and southern Namibia, western Zimbabwe, central Angola and the Congo basin represented by only Gabon (Figure 5).

**Figure 5.**
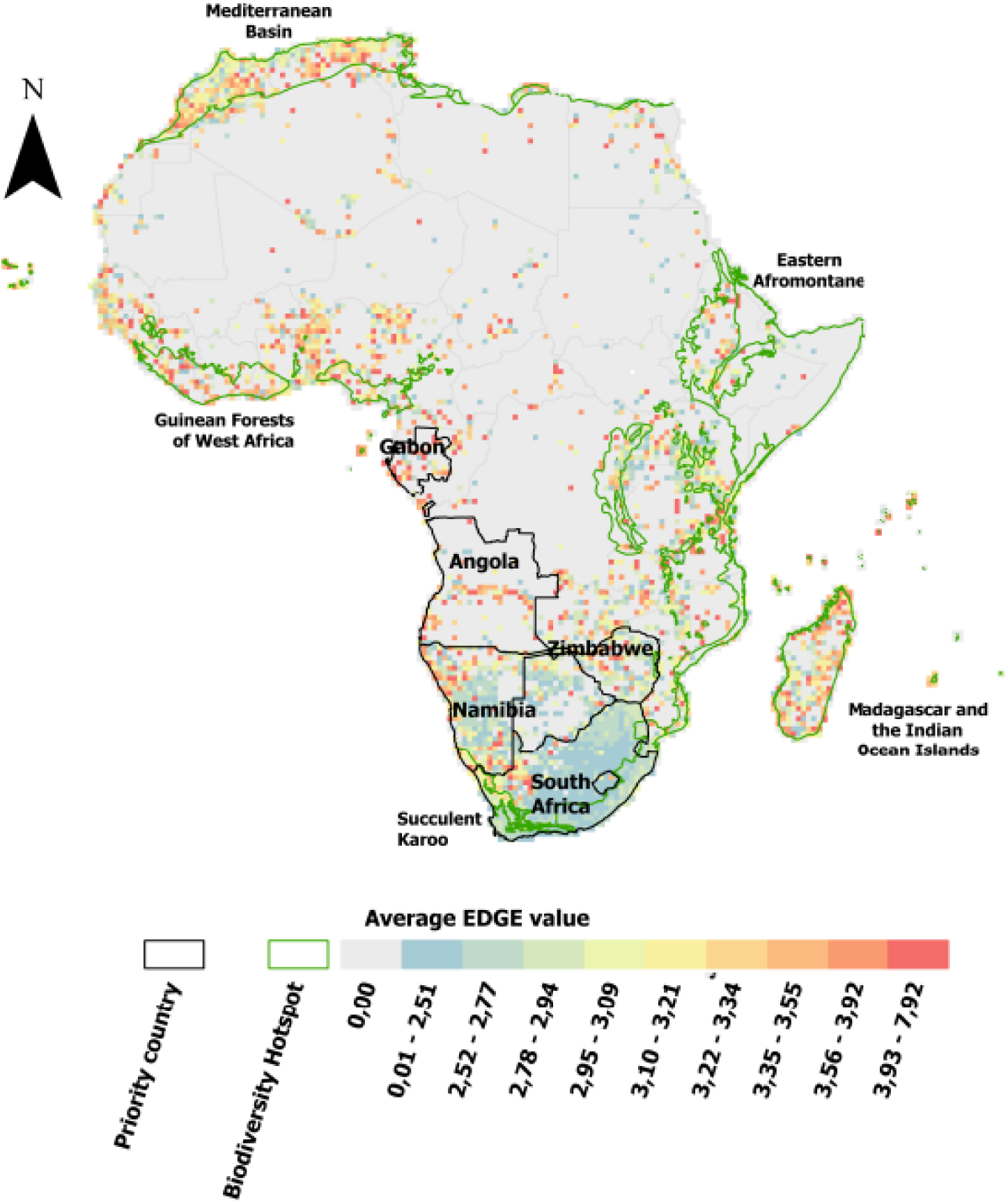
Geography of high-EDGE plant species in Africa

## 4. Discussion

### Our model performance is higher than that of most studies

We employed an automated deep-learning method to predict the unknown risk status of African plants. We used nine geographic features and climate features [27,58,62] which were used separately and in combination [27] We trained the neural network classification (nn-class) model on the five IUCN classes (LC, NT, VU, EN and CR). Our best performing model achieved a validation accuracy of 82.3%, higher than that reported, using similar techniques, for orchids (60%; [27]), trees (∼67%; [69]) and the vascular plants of South Africa (56.8%; [58]). Furthermore, the test accuracy of our model too is higher (67%) than that reported for the South Africa’s vascular plants (46%; Kandolo et al. [58]) and orchids (64%; [27]). These differences in model performances might be linked to differences in the algorithms and features used in different studies. For example, Zizka et al.’s [27] best model incorporated geographic and human footprint features; Silva et al. [69] utilized biome features whereas geographic and climate features were used in the present study. It is important to highlight that incorporating additional features (e.g., biological traits and phylogenetic features) did not seem to influence the optimal model performance [62]. Given the high performance of our model, we confidently used it to predict the extinction risk of the species with unknown conservation status and investigated the consequences of our predictions.

### Taxonomic and phylogenetic consequences of species extinction

Phylogenetic diversity (PD) as well as the evolutionarily distinct and globally endangered (EDGE) index are the two indicators recommended in the Convention on Biological Diversity’s Kunming-Montreal Global Biodiversity Framework to monitor the tree of life [82]. What do PD and EDGE indices say about the African tree of life?

Our analysis indicates that some families (e.g., Begoniaceae, Bruniaceae, Anacampserotaceae and Zamiaceae) are extinction-prone as they contain more threatened species than expected (but see Jetz et al. [83] for birds). The consequence of this pattern is that the loss of at-risk species would likely result in the loss of entire clades and therefore a great amount of evolutionary history. Indeed, we found that the amount of at-risk PD is far greater than expected, although threatened species are not particularly evolutionarily distinct. In one of our early studies, we showed through simulations that the loss of extinction-prone clades may drive the loss of a disproportionate number of branches from the tree of life (see [51]), resulting in a greater loss of PD than expected. Supporting this, our finding here indicates that, should all threatened species go extinct, we would lose 59% of all PD accumulated in the African plant tree of life. In the ongoing biodiversity crisis [20], which is forcing scientists and decision makers to strive towards bending the curve of biodiversity loss [5] and meet the ambitious goals to accelerate recovery, such loss is ill-afforded [83,84]. It is indeed ill-afforded given that each time species go extinct, branches of the tree-of-life are pruned, causing the loss of rare and unique species [85], as well as the loss of ‘feature diversity’ [53.86] which is required for ecosystem function and stability [46,52,55,56,87].

To avoid such loss, several studies call for the prioritization of high-ED [49, 82,83,88–93]. We explored whether high-ED species are more threatened than not. Although we found a significant link between ED and threat status, threatened species are not high-ED species. From a conservation perspective, this is an interesting finding since it implies that evolutionarily unique species are not particularly at risk. Most of the studies that showed interest into this relationship focused on vertebrates, e.g., mammals [94], birds [83], reptiles [93], and fish [43], and all reported no link between ED and threat status. On plants, however, we can recall only two studies that showed interest in testing the relationships between ED and threat status and both focused on gymnosperms where they showed that threatened species are high-ED species [49,95]. In the present study, we found that threatened African angiosperms are not high-ED species, a finding which, from a conservation perspective, is an interesting finding given that high-ED species have accumulated considerable evolutionary diversity [49,88,90,95].

### EDGE index and protection status of priority species

We employed the EDGE index, recommended to monitor the tree of life [6,82], to further question the African tree of life. Indeed, there is a global effort to inform conservation based on EDGE index (https://www.edgeofexistence.org/; [6]). We ranked species based on their EDGE score and found that *Ptisana robusta* (Marattiaceae), *Rhizoglossum bergianum* (Orchidaceae), *Angiopteris evecta* (Marattiaceae), *A. madagascariensis* (Marattiaceae) and *Philodendron hederaceum* (Oleaceae) lead the rank and become priority species for conservation. By providing such ranking for nearly 24000 plant species of Africa, our study complements the ongoing global effort to inform conservation based on EDGE.

Our analysis further showed that Madagascar, west African forests, Mediterranean basin, eastern Afromontane and part of some countries (Gabon, Angola, Namibia and South Africa) are priority regions for conservation because rich in high-EDGE species. It is important to highlight that the absence of the Congo basin (represented only by Gabon) does not mean the region is not of priority for conservation. It is likely because the Congo basin was not well represented in our sampling, failing to meet our criteria of at least 15 occurrence points for a species to be retained in our data. The Congo basin is politically instable, and this likely prevents repeated botanical expeditions causing poor sampling in GBIF. The absence of the Cape Floristic Region is also noteworthy. This is likely because most Cape clades are evolutionarily young [96,97], that is, not evolutionarily unique although highly threatened (see ref. [97]).

Unfortunately, we found that ∼72% of threatened species are outside protected areas and ∼79% of the top 1000-EDGE species are also not located in any form of protected areas. This tells there is still a long way to go to shorten the road to biodiversity recovery [20]. The effectiveness of the existing network of protected areas in protecting biodiversity is rarely assessed particularly in megadiverse countries ([98]; but see Kandolo et al. [58]). Our finding adds to the trend that existing networks in Africa is not enough to protect threatened taxa [19,99,100]. In this effort towards protection, since we found more abundant threatened species outside protected areas, we ought to (i) target species outside protected areas too in our risk assessment, (ii) adjust the network of protected areas in such a way that protects threatened species and (iii) to increase sampling efforts [19], particularly in rich but woefully under-explored regions such as the Congo basin.

It is now well established that at-risk species are clustered on a phylogeny (but see ref. [50] for cycads), and their loss might drive a disproportionate loss of the branches of the tree of life. Similarly, it is also well established that the loss of top ED species would drive a disproportionate loss of PD. Our study reveals that some clades are over-threatened (e.g., Begoniaceae, Bruniaceae, Anacampserotaceea, Zamiaceae) and the loss of threatened species would be equivalent to the loss of nearly 60% of the evolutionary history of the African flora. To halt this loss, we rank ∼24 000 species according to their EDGE score to inform targeted efforts for conservation. This is very important in the context of Africa where we found that current African network of protected areas do not cover high-EDGE species. Interestingly, some African biodiversity hotspots, e.g., west African forests, eastern Afromontane, Madagascar and mediterranean basin, harbour high-EDGE species. The management regime of these hotspots must be reinforced so that priority species can be preserved. Considering that vertebrates are central to most phylogenetically informed conservation studies and that there is a knowledge gap on this approach in Africa, our study contributes to closing such gap, reveals the extent of biodiversity crisis in Africa and points to the direction for recovery.

## Acknowledgements

We thank Mr Lucien Masime for his tremendous assistance with the Phyton analysis.

## SUPPLEMENTAL FIGURES

**Figure S1.** Sampling efforts as per available distribution data on GBIF for Africa

**Figure S2**. Performance of machine learning models for conservation assessments. (a) Accuracy and (b) loss trends for the 5-class conservation status model.

**Figure S3.** Confusion matrix of the maximal trained model of the detailed 5-class

**Figure S4.** Geography of threatened and high-EDGE species in relation to the network of protected areas in Africa. A) protection status of all threatened species and B) protection status of the top-1000-EDGE species.

## SUPPLEMENTAL TABLES

**Table S1.** Geoclimate data for DD species

**Table S2**. Geoclimate data for NE species

**Table S3**. Classification report on the 5-class prediction

**Table S4.** Residuals of the negative binomial model for each family

**Table S5**. ED and EDGE scores accompanied by threat status of African species

## References

1. IUCN. 2022 The IUCN Red List of threatened species v.2022–2. Gland, Switzerland: IUCN

2. IPBES. 2018 Summary for policymakers of the regional assessment report on biodiversity and ecosystem services for Africa of the Intergovernmental Science-Policy Platform on Biodiversity and Ecosystem Services. Archer E, Dziba LE, Mulongoy KJ, Maoela MA, Walters M, Biggs R, Cormier-Salem M-C, DeClerck F, Diaw MC, Dunham AE, Failler P, Gordon C, Harhash KA, Kasisi R, Kizito F, Nyingi WD, Oguge N, Osman-Elasha B, Stringer LC, Tito de Morais L, Assogbadjo A, Egoh BN, Halmy MW, Heubach K, Mensah A, Pereira L, Sitas N. (eds.). IPBES secretariat, Bonn, Germany. 49 pages.

3. Haddad NM, Brudvig LA, Clobert J, Davies KF, Gonzalez A, Holt RD, Lovejoy TE, Sexton JO, Austin MP, Collins CD, Cook WM, Damschen EI, Ewers RM, Foster BL, Jenkins CN, King AJ, Laurance WF, Levey DJ, Margules CR, Melbourne BA, Nicholls AO, Orrock JL, Song DX, Townshend JR. 2015 Habitat fragmentation and its lasting impact on Earth’s ecosystems. Sci. Adv. 20, e1500052. (doi:10.1126/sciadv.1500052)

4. Díaz S, Settele J, Brondízio ES, Ngo HT, Agard J, Arneth A, Balvanera P, Brauman KA, Butchart SHM, Chan KMA, Garibaldi LA, Ichii K, Liu J, Subramanian SM, Midgley GF, Miloslavich P, Molnár Z, Obura D, Pfaff A, Polasky S, Purvis A, Razzaque J, Reyers B, Chowdhury RR, Shin YJ, Visseren-Hamakers I, Willis KJ, Zayas CN. 2019 Pervasive human-driven decline of life on Earth points to the need for transformative change. Science 13, eaax3100. (doi:10.1126/science.aax3100)

5. Mace GM, Barrett M, Burgess ND, Cornell SE, Freeman R, Grooten M, Purvis A. 2018 Aiming higher to bend the curve of biodiversity loss. Nat. Sustain. 1, 448 – 451. (doi:10.1038/s41893-018-0130-0)

6. Gumbs R, Chaudhary A, Daru BH, Faith DP, Forest F, Gray CL, Kowalska A, Lee W-S, Pellens R, Pipins S, Pollock LJ, Rosindell J, Scherson RA, Owen NR. 2023 Indicators to monitor the status of the tree of life. Conserv. Biol. 37, e14138. (doi:10.1111/cobi.14138)

7. Milner-Gulland EJ, Addison P, Arlidge WN, Baker J, Booth H, Brooks T, Bull JW, Burgass MJ, Ekstrom J, zu Ermgassen SO, Fleming LV, Grub HM, von Hase A, Hoffmann M, Hutton J, JuffeBignoli D, ten Kate K, Kiesecker J, Kümpel NF, Maron M, Newing HS, Ole-Moiyoi K, Sinclair C, Sinclair S, Starkey M, Stuart SN, Tayleur C, Watson JE. 2021 Four steps for the Earth: mainstreaming the post-2020 global biodiversity framework. One Earth 4, 75–87. (doi:10.31235/osf.io/gjps6)

8. CBD. 2021 Kunming-montreal global biodiversity framework. Montreal, QC, Canada: CBD.

9. Millennium Ecosystem Assessment. 2005 Ecosystems and human well-being: biodiversity synthesis. World Resources Institute, Washington

10. Wake DB, Vredenburg VT. 2008 Are we in the midst of the sixth mass extinction? A view from the world of amphibians. Proc. Natl. Acad. Sci. U.S.A. 105, 11466–11473. (doi: 10.1073/pnas.0801921105)

11. Wilson EO. 1992 The diversity of life. Norton WW and Co, New York

12. IUCN. 2010 IUCN sampled red list index for plants. http://www.kew.org/science-conservation/kew-biodiversity/plants-at-risk/indexhtm.

13. Nic Lughadha E, Bachman SP, Leão TCC, Forest F, Halley JM, Moat J, Acedo C, Bacon KL, Brewer RFA, Gâteblé G, Gonçalves SC et al. 2020 Extinction risk and threats to plants and fungi. Plants People Planet 2, 389–408. (doi:10.1002/ppp3.10146)

14. Bar-On YM, Phillips R, Milo R. 2018 The biomass distribution on Earth. Proc. Natl. Acad. Sci. U.S.A. 115, 6506–6511. (doi:10.1073/pnas.1711842115)

15. Brummitt N, Araujo AC, Harris T. 2021 Areas of plant diversity—what do we know? Plants People Planet 3, 33–44. (doi:10.1002/ppp3.10110)

16. Meyer C, Weigelt P, Kreft H. 2016 Multidimensional biases, gaps and uncertainties in global plant occurrence information. Ecol. Lett. 19, 992–1006. (doi:10.1111/ele.12624)

17. Sharrock S, Hoft R, Dias BFS. 2018 An overview of recent progress in the implementation of the global strategy for plant conservation – a global perspective. Rodriguesia 69,1489–1511. (doi:10.1590/2175-7860201869401)

18. Bachman SP, Brown MJM, Leão TCC, Nic Lughadha E, Walker BE. 2024 Extinction risk predictions for the world’s flowering plants to support their conservation. New Phytol. 242, 797–808. (doi:10.1111/nph.19592)

19. Hoveka LN, van der Bank M, Bezeng BS, Davies TJ. 2020 Identifying biodiversity knowledge gaps for conserving South Africa’s endemic flora. Biodivers. Conserv. 29, 2803–2819. (doi:10.1007/s10531-020-01998-4)

20. Bezeng BS et al. 2025 An African perspective to biodiversity conservation in the twenty-first century. Phil. Trans. R. Soc. B 380, 20230443. (doi.org/10.1098/rstb.2023.0443)

21. Chapman CA et al. 2022 The future of sub-Saharan Africa’s biodiversity in the face of climate and societal change. Front. Ecol. Evol. 10, 790552. (doi:10.3389/fevo.2022.790552)

22. Africa Center for Strategic Studies. 2022 Rising sea levels besieging Africa’s booming coastal cities. See https://africacenter.org/wp-content/uploads/2023/02/Rising-Sea-Levels-ENG.pdf.

23. Stévart T, Dauby G, Lowry PP 2nd, Blach-Overgaard A, Droissart V, Harris DJ, Mackinder BA, Schatz GE, Sonké B, Sosef MSM, Svenning JC, Wieringa JJ, Couvreur TLP. 2019 A third of the tropical African flora is potentially threatened with extinction. Sci Adv. 20, eaax9444. (doi: 10.1126/sciadv.aax9444)

24. Chiaka JC, Liu G, Li H, Zhang W, Wu M, Huo Z, Gonella F. 2024 Land cover changes and management effectiveness of protected areas in tropical coastal area of sub-Saharan Africa. Environ. Sustain. Indic. 22, 100340. (doi:10.1016/j.indic.2024.100340)

25. Pollock LJ, O’Connor LMJ, Mokany K, Rosauer DF, Talluto L, Thuiller W. 2020 Protecting biodiversity (in all its complexity): New Models and Methods. Trends Ecol. Evol. 35,1119–1128. (doi: 10.1016/j.tree.2020.08.015)

26. Cooke R, Mancini F, Boyd RJ, Evans KL, Shaw A, Webb TJ, Isaac NJB. 2023 Protected areas support more species than unprotected areas in Great Britain but lose them equally rapidly. Biol. Conserv. 278,109884. (doi: 10.1016/j.biocon.2022.109884)

27. Zizka A, Silvestro D, Vitt P, Knight, TM. 2020 Automated conservation assessment of the orchid family with deep learning. Conserv. Biol. 35, 897–908. (doi:10.1111/cobi.13616)

28. Cowell CR, Lughadha EN, Anderson PML, Leão T, Williams J, Annecke WA. 2023 Prioritising species for monitoring in a South African protected area and the Red List for plants. Biodiv. Conserv. 32, 119–137. (doi:10.1007/s10531-022-02488-5)

29. FAO. 2018 A review of existing approaches and methods to assess climate change vulnerability of forests and forest-dependent people. Forestry Working Paper No. 5. Rome, FAO. 80 pp. Licence: CC BY-NC-SA 3.0 IGO.

30. Keith DA. 2009 The interpretation, assessment and conservation of ecological communities and ecosystems. Ecol. Manage. Restor. 10, S3–15. (doi:10.1111/j.1442-8903.2009.00453.x)

31. Barnosky AD, Matzke N, Tomiya S, Wogan GO, Swartz B, Quental TB, Marshall C, McGuire JL, Lindsey EL, Maguire KC, Mersey B, Ferrer EA. 2011 Has the Earth’s sixth mass extinction already arrived? Nature 471, 51–7. (doi:10.1038/nature09678)

32. Ceballos G, Ehrlich PR, Barnosky AD, García A, Pringle RM, Palmer TM. 2015 Accelerated modern human-induced species losses: Entering the sixth mass extinction. Sci Adv. 19, e1400253. (doi: 10.1126/sciadv.1400253)

33. IUCN. 2024 The IUCN red list of threatened species. Version 2023–1. See https://www.iucnredlist.org.

34. Brummitt NA, Bachman SP, Griffiths-Lee J, Lutz M, Moat JF, Farjon A, Donaldson JS, Hilton-Taylor C, Meagher TR, Albuquerque S et al. 2015 Green plants in the red: a baseline global assessment for the IUCN sampled red list index for plants. PLoS One 10, e0135152. (doi:10.1371/journal.pone.0135152)

35. Raimondo D, von Staden L, Foden W, Victor J, Helme N, Turner R, Kamundi D, Manyam P. 2009 Red list of South African plants. Pretoria, South Africa: South African National Biodiversity Institute (SANBI Publishing

36. Dinerstein E, Vynne C, Sala E, Joshi AR, Fernando S, Lovejoy TE, Mayorga J, Olson D, Asner GP, Baillie JEM et al. 2019. A global deal for nature: guiding principles, milestones, and targets. Sci Adv. 5, eaaw2869. (doi: 10.1126/sciadv.aaw2869)

37. Maxwell SL, Cazalis V, Dudley N, Hoffmann M, Rodrigues ASL, Stolton S, Visconti P, Woodley S, Kingston N, Lewis E et al. 2020 Area-based conservation in the twenty-first century. Nature 586, 217–227. (doi:10.1038/s41586-020-2773-z)

38. Walker BE, Leão TCC, Bachman SP, Lucas E, Lughadha EN. 2021 Evidence-based guidelines for developing automated conservation assessment methods. Conserv Biol. 37, e13992. (doi: 10.1111/cobi.13992)

39. Cazalis V, Di Marco M, Butchart SHM, Akçakaya HR, González-Suárez M, Meyer C, Clausnitzer V, Böhm M, Zizka A, Cardoso P, Schipper AM, Bachman SP, Young BE, Hoffmann M, Benítez-López A, Lucas PM, Pettorelli N, Patoine G, Pacifici M, Jörger-Hickfang T, Brooks TM, Rondinini C, Hill SLL, Visconti P, Santini L. 2022 Bridging the research-implementation gap in IUCN Red List assessments. Trends Ecol Evol. 37, 359–370. (doi: 10.1016/j.tree.2021.12.002)

40. Botanic Gardens Conservation International. 2021 State of the World’s Trees. Richmond, UK: Botanic Gardens Conservation International

41. Bachman S, Moat J, Hill A, de la Torre J, Scott B. 2011 Supporting Red List threat assessments with GeoCAT: geospatial conservation assessment tool. ZooKeys 150, 117–126. (doi: 10.3897/zookeys.150.2109)

42. Moat J. 2020. rCAT: conservation assessment tools, version: 0.1.6. [WWW document] URL https://CRAN.R-project.org/package=rCAT

43. Bachman SP, Brown MJM, Leão TCC, Nic Lughadha E, Walker BE. 2024 Extinction risk predictions for the world’s flowering plants to support their conservation. New Phytol. 242, 797–808. (doi: 10.1111/nph.19592)

44. Adeoba M, Tesfamichael SG, Yessoufou K. 2019 Preserving the tree of life of the fish family Cyprinidae in Africa in the face of the ongoing extinction crisis 1. Genome 62, 170–182. (doi: 10.1139/gen-2018-0023)

45. Tonini JFR. Beard, KH, Ferreira RB, Jetz W, Pyron RA. 2016 Fully-sampled phylogenies of squamates reveal evolutionary patterns in threat status. Biol. Conserv. 204, 23–31. (doi:10.1016/j.biocon.2016.03.039)

46. Veron S, Davies TJ, Cadotte MW, Clergeau P, Pavoine S. 2017 Predicting loss of evolutionary history: Where are we? Biol. Rev. 92, 271–291. (doi: 10.1111/brv.12228)

47. Faith DP. 2008 Threatened species and the potential loss of phylogenetic diversity: Conservation scenarios based on estimated extinction probabilities and phylogenetic risk analysis. Conserv. Biol. 22, 1461–1470. (doi: 10.1111/j.1523-1739.2008.01068.x.)

48. Mooers A.Ø, Faith DP, Maddison WP 2008 Converting endangered species categories to probabilities of extinction for phylogenetic conservation prioritization. PLoS One 3, e3700. (doi:10.1371/journal.pone.0003700)

49. Thuiller W, Lavergne, S, Roquet C, Boulangeat I, Lafourcade B Araujo MB. 2011 Consequences of climate change on the tree of life in Europe. Nature 470, 531– 534. (doi:10.1038/nature09705)

50. Yessoufou K, Daru BH, Tafirei, R., Elansary, H.O, Rampedi I. 2017. Integrating biogeography, threat and evolutionary data to explore extinction crisis in the taxonomic group of cycads. Ecol. Evol. 7, 2735–2746. (doi: 10.1002/ece3.2660)

51. Davies TJ, Yessoufou K. 2013 Revisiting the impacts of non-random extinction on the tree-of-life. Biol. Lett. 9, 20130343. (doi:10.1098/rsbl.2013.0343)

52. Forest F, Grenyer R, Rouget M, Davies TJ, Cowling RM, Faith DP, Balmford A, Manning JC, Procheş S, van der Bank M, Reeves G, Hedderson TA, Savolainen V. 2007 Preserving the evolutionary potential of floras in biodiversity hotspots. Nature 445, 757–760. (doi: 10.1038/nature05587)

53. Faith, DP. 1992 Conservation evaluation and phylogenetic diversity. Biol. Conserv. 61, 1–10. (doi:10.1016/0006-3207(92)91201-3)

54. Winter M, Devictor V, Schweiger O. 2012. Phylogenetic diversity and nature conservation: where are we? Trends Ecol. Evol. 28, 199–204. (doi:10.1016/j.tree.2012.10.015)

55. Faith DP, Magallün S, Hendry AP, Conti E, Yahara T, Donoghue MJ. 2010 Ecosystem services: an evolutionary perspective on the links between biodiversity and human well-being. Curr. Opin. Environ. Sustain. 2, 66–74. (doi:10.1016/j.cosust.2010.04.002)

56. Faith DP, Pollock LJ. 2014 Phylogenetic diversity and the sustainable use of biodiversity. In Applied ecology and human dimensions in biological conservation. Edited by Lyra-Jorge MC and Verdade LM. Springer, Berlin, Heidelberg. pp.35–52. (doi:10.1007/978-3-642-54751-5_3)

57. Zizka A, Andermann T, Silvestro D. 2021 IUCNN—Deep learning \approaches to approximate species’ extinction risk. Divers. Distrib. 28, 227–241. (doi.org/10.1111/ddi.13450)

58. Kandolo BS, Yessoufou K, Kganyago M. 2024 Effectiveness of South Africa’s network of protected areas: Unassessed vascular plants predicted to be threatened using deep neural networks are all located in protected areas. Ecol Evol. 14, e70229. (doi: 10.1002/ece3.70229)

59. Deneu B, Servajean M, Bonnet P, Botella C, Munoz F, Joly A. 2021 Convolutional neural networks improve species distribution modelling by capturing the spatial structure of the environment. PLoS Comput Biol. 17, e1008856. (doi: 10.1371/journal.pcbi.1008856)

60. Tuia D. 2022 Kellenberger, B., Beery, S. et al. Perspectives in machine learning for wildlife conservation. Nat Commun 13, 792. (doi:10.1038/s41467-022-27980-y)

61. Taherdoost, H. 2023. Deep Learning and Neural Networks: Decision-Making Implications. Symmetry 15, 1723. (doi:10.3390/sym15091723)

62. Chen J, Ding C, He D, Ding L, Ji S, Du T, Sun J, Huang M, Tao J. 2023. Assessing the conservation status of Chinese freshwater fish using deep learning. Rev. Fish Biol. Fish. 33, 1505–1521. (doi:10.1007/s11160-023-09792-5)

63. Yessoufou K, Daru BH, Davies TJ. 2012 Phylogenetic Patterns of Extinction Risk in the Eastern Arc Ecosystems, an African Biodiversity Hotspot. PLOS One 7, e47082. (doi:10.1371/journal.pone.0047082)

64. Tucker CM, Cadotte MW, Davies TJ, Rebelo TG. 2012 Incorporating geographical and evolutionary rarity into conservation prioritization. Conserv. Biol. 26, 593–601. (doi: 10.1111/j.1523-1739.2012.01845.x)

65. Cadotte MW, Dinnage R, Tilman D. 2012 Phylogenetic diversity promotes ecosystem stability. Ecology 93, S223–S233. (doi:10.1890/11-0426.1)

66. GBIF.org (25 October 2024) GBIF Occurrence Download 10.15468/dl.auj28h66.

67. South African National Biodiversity Institute (SANBI). (2024). Statistics: Red list of South African plants. Redlist.sanbi.Org

68. Maldonado C, Molina CI, Zizka A, Persson C, Taylor CM, Albán J, Chilquillo E, Rønsted N, Antonelli A. 2015 Estimating species diversity and distribution in the era of Big Data: To what extent can we trust public databases? Glob. Ecol. Biogeogr. 24, 973–984. (doi:10.1111/geb.12326)

69. Silva SV, Andermann T, Zizka A, Kozlowski G and Silvestro D (2022) Global Estimation and Mapping of the Conservation Status of Tree Species Using Artificial Intelligence. Front. Plant Sci. 13, 839792. (doi: 10.3389/fpls.2022.839792)

70. Rivers MC, Taylor L, Brummitt NA, Meagher TR, Roberts DL, Lughadha EN. 2011 How many herbarium specimens are needed to detect threatened species? Biol. Conserv. 144, 2541–2547. (doi:10.1016/j.biocon.2011.07.014)

71. Zizka A, Silvestro D, Vitt P, Knight TM 2021 Automated conservation assessment of the orchid family with deep learning. Conserv Biol 35, 897–908. (doi:10.1111/cobi.13616)

72. Zizka A, Andermann T, Silvestro D. 2022 IUCNN—Deep learning approaches to approximate species’ extinction risk. Divers. Distrib. 28, 227–241. (doi:10.1111/ddi.13450)

73. Karger D, Conrad O, Böhner J, Kawohl T, Kreft H, Soria-Auza RW, Zimmermann NE, Linder HP, Kessler M. 2017 Climatologies at high resolution for the earth’s land surface areas. Sci Data 4, 170122. (doi:10.1038/sdata.2017.122)

74. Pedregosa F, Varoquaux G, Gramfort A, Michel V, Thirion B, Grisel O, Blondel M, Prettenhofer P, Weiss, R, Dubourg V, Vanderplas J. Passos A, Cournapeau D, Brucher M, Perrot M, Duchesnay E. 2011 Scikit-learn: Machine learning in Python. J. Mach. Learn. Res. 12, 2825−2830. (doi:10.48550/arXiv.1201.0490)

75. Kingma DP, Ba J. 2015 Adam: A Method for Stochastic Optimization. Conference Paper at the 3rd International Conference for Learning Representations, San Diego. arXiv:1412.6980ss

76. Chawla NV, Bowyer KW, Hall LO, Kegelmeyer WP. 2002 SMOTE: synthetic minority over-sampling technique. J. Artif. Intell. Res 16, 321–357. (doi:10.48550/arXiv.1106.1813.)

77. Gal Y, Ghahramani Z. 2016 Dropout as a Bayesian approximation: Representing model uncertainty in deep learning. International Conference on Machine Learning.

78. Jin Y, Qian HV. 2019 PhyloMaker: An R package that can generate very large phylogenies for vascular plants. Ecography 42, 1353–1359. (doi.org/10.1111/ecog.04434)

79. Smith SA, Brown JW. 2018 Constructing a broadly inclusive seed plant phylogeny. Am. J. Bot. 105, 302–314. (doi: 10.1002/ajb2.1019)

80. Zanne AE, Tank DC, Cornwell WK, Eastman JM, Smith SA, FitzJohn RG, McGlinn DJ, O’meara BC, Moles AT, Reich PB, et al. 2014 Three keys to the radiation of angiosperms into freezing environments. Nature 506, 89–92. (doi: 10.1038/nature12872.)

81. Isaac NJB, Turvey ST, Collen B, Waterman C, Baillie JEM. 2007 Mammals on the EDGE: Conservation priorities based on threat and phylogeny. PLoS One, 2, e296. (doi:10.1371/journal.pone.0000296)

82. Secretariat of the Convention on Biological Diversity. 2022 Monitoring framework for the Kunming-Montreal global biodiversity framework. https://www.cbd.int/doc/c/179e/aecb/592f67904bf07dca7d0971da/cop-15-l-26-en.pdf

83. Jetz W, Rahbek C, Colwell RK. 2004 The coincidence of rarity and richness and the potential signature of history in centres of endemism. Ecol. Lett. 7, 1180–1191. (doi:10.1111/j.1461-0248.2004.00678.x)

84. Pellens R, Grandcolas P. 2016 Phylogenetics and conservation biology: drawing a path into the diversity of life. In Biodiversity conservation and phylogenetic systematics. Springer International Publishing. pp. 1–15.

85. Winter M, Devictor V, Schweiger O. 2012 Phylogenetic diversity and nature conservation: where are we? Trends Ecol. Evol. 28, 199–204. (doi:10.1016/j.tree.2012.10.015)

86. Crozier RH. 1997 Preserving the information content of species: genetic diversity, phylogeny and conservation worth. Annu. Rev. Ecol. Syst. 28, 243 – 268. (doi:10.1146/annurev.ecolsys.28.1.243)

87. Cadotte MW, Cardinale BJ, Oakley TH. 2008 Evolutionary history and the effect of biodiversity on plant productivity. Proc. Natl Acad. Sci. USA 105, 17 012 – 17 017. (doi:10.1073/pnas.080596210)

88. Redding DW, Hartmann K, Mimoto A, Bokal D, Devos M, Mooers A. 2008 Evolutionarily distinctive species often capture more phylogenetic diversity than expected. J. Theor. Biol. 251, 606–615. (doi:10.1016/j.jtbi.2007.12.006)

89. Redding DW, DeWolff CV, Mooers AO. 2010 Evolutionary distinctiveness, threat status, and ecological oddity in primates. Conserv. Biol. 24, 1052–1058. (doi:10.1111/j.1523-1739.2010.01532.x)

90. Redding DW, Mooers AO, Sekercioglu CH, Collen B. 2015 Global evolutionary isolation measures can capture key local conservation species in Nearctic and Neotropical bird communities. Philos. Trans. R. Soc. B Biol. Sci.B 370, 20140013. (doi:10.1098/rstb.2014.0013)

91. Warren WC, Hillier LW, Graves JAM, Birney E, Ponting CP, Grutzner F et al. 2008 Genome analysis of the platypus reveals unique signatures of evolution. Nature 453, 175–183. (doi:10.1038/nature06936)

92. Daru BH. Yessoufou K. Mankga LT, Davies TJ. 2013 A global trend towards the loss of evolutionarily unique species in mangrove ecosystems. PLoS One 8, e66686. (doi:10.1371/journal.pone.0066686)

93. Tonini JFR, Beard KH, Ferreira RB, Jetz W, Pyron RA. 2016 Fully-sampled phylogenies of squamates reveal evolutionary patterns in threat status. Biol. Conserv. 204, 23–31. (doi:10.1016/j.biocon.2016.03.039)

94. Arregoitia LD, Blomberg SP, Fisher DO. 2013 Phylogenetic correlates of extinction risk in mammals: species in older lineages are not at greater risk. Proc. R. Soc. B Biol. Sci. 280, 20131092. (doi:10.1098/rspb.2013.1092)

95. Forest Forest F, Moat J, Baloch E, Brummitt NA, Bachman SP, Ickert-Bond S, Hollingsworth PM, Liston A, Little DP, Mathews S, Rai H, Rydin C, Stevenson DW, Thomas P, Buerki S. 2018 Gymnosperms on the EDGE. Sci Rep. 8, 6053. (doi:10.1038/s41598-018-24365-4)

96. Verboom GA, Jenny K. Archibald, Freek T. Bakker, Dirk U. Bellstedt, Ferozah Conrad, Leanne L. Dreyer, Félix Forest, Chloé Galley, Peter Goldblatt, Jack F. Henning, Klaus Mummenhoff, H. Peter Linder, A. Muthama Muasya, Kenneth C. Oberlander, Vincent Savolainen, Deidre A. Snijman, Timotheüs van der Niet, Tracey L. Nowell, 2009. Origin and diversification of the Greater Cape flora: Ancient species repository, hot-bed of recent radiation, or both? Mol. Phylogenet. Evol. 51, 44–53. (doi:10.1016/j.ympev.2008.01.037)

97. Davies TJ et al. 2011 Extinction risk and diversification are linked in a plant biodiversity hotspot. PLoS Biol. 9, e1000620. (doi:10.1371/journal.pbio.1000620)

98. Paknia O, Rajaei ShH, Koch A. 2015 Lack of well-maintained natural history collections and taxonomists in megadiverse developing countries hampers global biodiversity exploration. Org Divers Evol 15, 619–629. (doi:10.1007/s13127-015-0202-1)

99. Simaika JP, Samways MJ. 2009 Reserve selection using red listed taxa in three global biodiversity hotspots: Dragonflies in South Africa. Biol. Conserv. 142, 638–651. (doi:10.1016/j.biocon.2008.11.012)

100. Tolley KA, Weeber J, Maritz B, Verburgt L, Bates MF, Conradie W, Hofmeyr MD, Turner AA, da Silva JM, Alexander GJ. 2019 No safe haven: Protection levels show imperilled South African reptiles not sufficiently safe-guarded despite low average extinction risk. Biol. Conserv 233, 61–72. (doi:10.1016/j.biocon.2019.02.006)

101. Tiawoun MAP, Phogole B, Kandolo BS, Yessoufou K 2025. Over half of the phylogenetic diversity accumulated on the African plant tree of life may be eroded under current biodiversity crisis even though threatened species are not evolutionarily unique. Dataset, DOI: 10.6084/m9.figshare.28390718

