## Supplementary material for "Over half of the phylogenetic diversity accumulated on the African plant tree of life may be eroded under current biodiversity crisis even though threatened species are not evolutionarily unique": Phyton code.txt

import os
import numpy as np
import pandas as pd
import tensorflow as tf
from tensorflow import keras
from tensorflow.keras import layers
from tensorflow.keras.regularizers import l2
from sklearn.model_selection import train_test_split
from sklearn.metrics import classification_report, confusion_matrix
from sklearn.preprocessing import StandardScaler, OneHotEncoder
import matplotlib.pyplot as plt
import seaborn as sns
from imblearn.over_sampling import SMOTE  # Import SMOTE

np.set_printoptions(suppress=True, precision=1)

# Set seeds for reproducibility
np.random.seed(42)
tf.random.set_seed(42)


# Set working directory (adjust this path)
wd = "C:/Users/LUC MAS/Desktop/Appendices/csv files/true data/Features"

# ---Load the Data
train_feat_file = "geoclimate1.csv"
train_label_file = "AfricaLabels1.csv"

features = pd.read_csv(os.path.join(wd, train_feat_file), delimiter=";", decimal=",").to_numpy()
labels = pd.read_csv(os.path.join(wd, train_label_file), delimiter=";", decimal=",").to_numpy()


#Convert labels to integers (0: LC, 1: NT, 2: VU, 3: EN, 4: CR)
label_mapping = {
   "LC": 0,
   "NT": 1,
   "VU": 2,
   "EN": 3,
   "CR": 4
}
integer_labels = np.vectorize(label_mapping.get)(labels[:, 1])

# One-hot encoding of the labels
onehot_encoder = OneHotEncoder(sparse_output=False)
onehot_labels = onehot_encoder.fit_transform(integer_labels.reshape(-1, 1))

# Convert features to float
Xt = features[:, 1:].astype(float)

# Normalization
scaler = StandardScaler()
Xt_normalized = scaler.fit_transform(Xt)

# ---Split the data into training and testing sets
X_train, X_test, y_train, y_test = train_test_split(Xt_normalized, onehot_labels, test_size=0.2, random_state=42)

# Step 4: Apply SMOTE to address class imbalance
smote = SMOTE(random_state=42)
X_train_resampled, y_train_resampled = smote.fit_resample(X_train, y_train)

# Step 4: Build the Model with L2 Regularization
def build_multiclass_classification_model(input_shape, regularization_rate=0.03): 
    model = keras.Sequential()
    model.add(layers.Dense(71, activation='relu', input_shape=(input_shape,), kernel_regularizer=l2(regularization_rate))) 
    model.add(layers.Dense(46, activation='relu', kernel_regularizer=l2(regularization_rate))) 
    model.add(layers.Dense(5, activation='softmax'))  # Softmax for multi-class classification
    
    model.compile(optimizer='adam', loss='categorical_crossentropy', metrics=['accuracy'])
    return model

# Create the model
model = build_multiclass_classification_model(X_train_resampled.shape[1])
model.summary()

# Step 5: Train the Model
history = model.fit(X_train_resampled, y_train_resampled, epochs=175, validation_split=0.13, batch_size=66, verbose=1) 

# ---Make Predictions
y_pred = model.predict(X_test)
y_pred_classes = np.argmax(y_pred, axis=1)  # Convert probabilities to class predictions
y_test_classes = np.argmax(y_test, axis=1)  # Convert one-hot encoded labels back to class labels

# ---Evaluate the Model
print("Classification Report:\n", classification_report(y_test_classes, y_pred_classes))
conf_matrix = confusion_matrix(y_test_classes, y_pred_classes)
print(conf_matrix)

# ---Visualize the confusion matrix
plt.figure(figsize=(10, 7))
sns.heatmap(conf_matrix, annot=True, fmt='d', cmap='Blues', 
            xticklabels=["LC", "NT", "VU", "EN", "CR"], 
            yticklabels=["LC", "NT", "VU", "EN", "CR"])
plt.title('Confusion Matrix')
plt.xlabel('Predicted Label')
plt.ylabel('True Label')
plt.show()

# Function to plot learning curve
def plot_learning_curve(history):
    plt.figure(figsize=(12, 5))

    # Plot accuracy
    plt.subplot(1, 2, 1)
    plt.plot(history.history['accuracy'], label='Training Accuracy')
    plt.plot(history.history['val_accuracy'], label='Validation Accuracy')
    plt.title('Accuracy over Epochs')
    plt.xlabel('Epochs')
    plt.ylabel('Accuracy')
    plt.legend()

    # Plot loss
    plt.subplot(1, 2, 2)
    plt.plot(history.history['loss'], label='Training Loss')
    plt.plot(history.history['val_loss'], label='Validation Loss')
    plt.title('Loss over Epochs')
    plt.xlabel('Epochs')
    plt.ylabel('Loss')
    plt.legend()

    plt.tight_layout()
    plt.show()

# ---Plot Learning Curve
plot_learning_curve(history)

# ---Load Unlabeled Data and Make Predictions
unlabeled_data_file = "featuresDD.csv"  # Replace with your actual file path
unlabeled_features = pd.read_csv(os.path.join(wd, unlabeled_data_file), delimiter=";", decimal=",").to_numpy()

# Extract the features and species names for the unlabeled data
species = unlabeled_features[:, 0].astype(str)  # Adjust this if the first column isn't the species name

# If the features matrix has an extra column (like an ID column), you can slice out only the feature columns.
# For example, if the first column is an ID, you can ignore it like this:
unlabeled_features = unlabeled_features[:, 1:]  # This removes the first column (index/ID column)

# Normalize the features using the same scaler as the training data
unlabeled_features_normalized = scaler.transform(unlabeled_features.astype(float))

# Predict using the trained model
unlabeled_predictions = model.predict(unlabeled_features_normalized)
unlabeled_pred_classes = np.argmax(unlabeled_predictions, axis=1)


# ---Plot Histogram of Predictions (Multicolor) ---
# Define class-to-color mapping (light to dark)
color_map = {
    0: ['#98FB98', '#006400'],  # LC: light green to dark green
    1: ['#D2B48C', '#8B4513'],  # NT: light brown to dark brown
    2: ['#FFFF00', '#FFD700'],  # VU: light yellow to dark yellow
    3: ['#FFA500', '#FF4500'],  # EN: light orange to dark orange
    4: ['#FF6347', '#8B0000']   # CR: light red to dark red
}

# Frequency of predictions for each class (0 to 4)
class_counts = np.bincount(unlabeled_pred_classes, minlength=5)

# Create the histogram with separate bars for each class and increasing darkness based on frequency
plt.figure(figsize=(10, 6))

# Bar width and positioning
bar_width = 0.9  # Adjust bar width for spacing between classes
positions = np.arange(5)  # Class positions

# Loop through each class and plot bars with corresponding color
for i, count in enumerate(class_counts):
    # Use the gradient from light to dark for each class based on its frequency
    color = color_map[i][0] if count < max(class_counts) // 2 else color_map[i][1]  # Light if count is lower, dark if higher
    plt.bar(positions[i], count, width=bar_width, color=color, label=f"Class {i}" if count > 0 else "")

# Add labels and title
plt.title("Number of species per IUCNN category")
plt.xlabel("Class Label")
plt.ylabel("Frequency")
plt.xticks(positions, ['LC', 'NT', 'VU', 'EN', 'CR'])  # Label classes
plt.legend(title="Classes", loc="upper right")

# Adjust layout to avoid overlap
plt.tight_layout()

# Show the plot
plt.show()


# Get the predicted probabilities for each class (LC, NT, VU, EN, CR)
unlabeled_pred_probs = model.predict(unlabeled_features_normalized)

# Get the predicted labels (class indices) by taking the argmax of the probabilities
unlabeled_pred_classes = np.argmax(unlabeled_pred_probs, axis=1)


# Create a DataFrame with only the species and prediction results
prediction_df = pd.DataFrame({
    'species': species,
    'P(LC)': unlabeled_pred_probs[:, 0],
    'P(NT)': unlabeled_pred_probs[:, 1],
    'P(VU)': unlabeled_pred_probs[:, 2],
    'P(EN)': unlabeled_pred_probs[:, 3],
    'P(CR)': unlabeled_pred_probs[:, 4],
    'Predicted_Label': np.array(["LC", "NT", "VU", "EN", "CR"])[unlabeled_pred_classes]
})

# ---Save to CSV and Excel
prediction_df.to_csv(os.path.join(wd,'Unlabeled_Data_Predictions5_rec.csv'))
prediction_df.to_excel(os.path.join(wd,'Unlabeled_Data_Predictions5_rec.xlsx'))

# Print a confirmation message
print("Predictions saved to 'Unlabeled_Data_Predictions.csv' and 'Unlabeled_Data_Predictions.xlsx'.")
