## Supplementary material for "Over half of the phylogenetic diversity accumulated on the African plant tree of life may be eroded under current biodiversity crisis even though threatened species are not evolutionarily unique": R script_Africa_Extinction risk.R

```
#analysis of extinction risk in Africa
#by Prof. Kowiyou Yessoufou

rm(list=ls(all=TRUE))

# predicting IUCN status of DD and NE species

#IUCNN installation
install.packages("devtools")
library(usethis)
library(devtools)
install.packages(IUCNN)
library(IUCNN)

#Loading data
## Training occurrences
#choose.files()
train_occ <- read.table("C:\\Users\\sam\\Desktop\\occ.txt",header=TRUE)
attach(train_occ)
names(train_occ)

## Training labels: Detailed classification
Choose.files()
train_lab <- read.table("C:\\Users\\sam\\Desktop\\Lab.txt",header=TRUE)

## Training labels: Binary classification labels
#Choose.files()
train_lab2 <- read.table("C:\\Users\\sam\\Desktop\\broadlabels.txt",header=TRUE)


# Features generation
## Geographic features
geo_feat <-iucnn_prepare_features(train_occ, type="geographic",impute_features=FALSE)

## Climate features
climate_feat <-iucnn_prepare_features(train_occ, type="climate",impute_features=FALSE)

## Geographic + climate
allfeat <-iucnn_prepare_features(train_occ,type=c("geographic","climate"),impute_features=FALSE)


#Labels preparation
##For the 5-class

labfive <- iucnn_prepare_labels(train_lab,geo_feat,level="detail")

#
#### From here we moved to Phyton analysis. See Phyton code attached in separate file


#phylogenetic and taxonomic data analysis
# load the package
library(ape)
library("V.PhyloMaker")
library(utf8)

# input the list of all woody species recorded in all plots
species_list <- read.table("C:\\Users\\kowiyouy\\OneDrive - University of Johannesburg\\Desktop\\UJ\\postdoc\\2024\\Andre\\papers\\paper 1_iucn\\data\\final\\species_list_africa.txt",header=TRUE)
# generate the phylogeny presented in
#species_list$species
#utf8_normalize(species_list$species)
#as_utf8(species_list$species, normalize = TRUE)
#as_utf8(species_list$species)
#utf8_valid(species_list$species)

tree.africa <- phylo.maker(sp.list = species_list,
                           tree = GBOTB.extended, nodes = nodes.info.1, scenarios="S3")
tree.africa <- tree.africa$scenario.3

write.nexus(tree.africa, file = "tree_africa")


plot(tree.africa,cex=0.001)

# call the tree
library(caper)
tree_africa<-read.nexus("C:\\Users\\kowiyouy\\OneDrive - University of Johannesburg\\Desktop\\UJ\\postdoc\\2024\\Andre\\papers\\paper 1_iucn\\data\\tree_africa.txt")

# calculate ED (evolutionary distinctiveness)
ed_africa <- ed.calc(tree_africa)
ed_africa1 <- ed_africa$spp

#save ED values
write.table(ed_africa1, file = "ed_africa2")

# Analysis of extinction risk
#question 1: Are threatened species evolutionary more distinct than not?

data_ext <- read.table("C:\\Users\\kowiyouy\\OneDrive - University of Johannesburg\\Desktop\\UJ\\postdoc\\2024\\Andre\\papers\\paper 1_iucn\\data\\final\\data_ext_africa.txt",header = TRUE)
attach(data_ext)
names(data_ext)

# dividing the plot window into three plot panels
par(mfrow=c(1,2))

#fitting models and graphically represent the relationships between ED and IUCN status
#model 1: looking at iucn individually

model1 <- aov(log(ED)~iucn_predicted_transf,data=data_ext)
summary(model1)
#                          Df  Sum Sq MeanSq  Fvalue Pr(>F)    
#iucn_predicted_transf     4   3099   774.6   366.3   <2e-16 ***
#  Residuals            23328  49333  2.1  

boxplot(log(ED)~iucn_predicted_transf,data=data_ext,main="A",notch=TRUE,xlab="IUCN categories",ylab="Evolutionary distinctiveness (log)",col=c("blue","green","yellow","orange","red"), names=c("LC","NT","VU","EN","CR"))

#model 2: combining iucn status into threatened vs nonthreatened

model2 <- aov(log(ED)~iucn_threat_status_binary_cat,data=data_ext)
summary(model2)
#                                Df    Sum Sq  Mean Sq Fvalue Pr(>F)    
#iucn_threat_status_binary_cat    1    2105    2104.6   975.6 <2e-16 ***
#  Residuals                     23331 50327   2.2  
boxplot(log(ED)~iucn_threat_status_binary_cat,data=data_ext,main="B",notch=TRUE,xlab="threat categories",ylab="",col=c("blue","red"))

#### Question 2: Are threatened species clustered on the phylogeny? 
#Testing phylogenetic signal in extinction risk using the D statistics (Fritz & Purvis 2010)

tree_africa$node.label<-NULL

africa_ext_data <- comparative.data(tree_africa, data_ext, species)
PhyloD_iucn <- phylo.d(africa_ext_data, binvar=iucn_threat_status_binary_num)
print(PhyloD_iucn)
plot(PhyloD_iucn,main="threat status")


###Question 3: Are we going to lose more PD than at random if all all threatened species go extinct?

# Calculate total PD on the phylogenetic tree
total_pd <- sum(tree_africa$edge.length)

#form the list of all threatened species

#firstly, create the list of nonthreatened species
list_nonthreatened <- data_ext$species[!data_ext$iucn_threat_status_binary_cat=="threatened"]

#secondly, drop all nonthreatened species from the tree to have a tree of only threatened species
extinct.tree <- drop.tip(tree_africa,as.vector(list_nonthreatened))

# Calculate threatened PD i.e. total PD on the tree containing only threatened species
pd_threatened <- sum(extinct.tree$edge.length)

#Calculate the proportion of threatened pd

100*pd_threatened/total_pd # ~ 58.64629 = 59%

#Investigating how a random loss of a number of species equivalent to the number of threatened species would impact the tree
# This requires the following simulation of species loss (same number of species as real number of threatened species are pruned randomly 1000 times from the tree)

# calculate number of threatened species on the phylogeny
number_threatened_species <- length(extinct.tree$tip.label)

#number_threatened_species= 12647

# simulation of loss of 12647 species pruned randomly 1,000 times from the tree, and calculation of pd remaining on the tree each time after a random loss of 12647 species
pd <- NULL
for(x in 1:1000){
  rand.tree <- drop.tip(tree_africa,sample(tree_africa$tip.label,number_threatened_species))
  pd[x] <- sum(rand.tree$edge.length)
}

# Calculation of the pd corresponding to each set of 12647 species randomly lost from the tree
pd_random_loss <- total_pd-pd # there are 1,000 values of pd_random_loss

#plot the results in a form of histogram
par(mfrow=c(1,1))

hist(total_pd-pd,main="",xlim = c(130000,185000),xlab="randomly lost pd") # histogram of random loss
abline(v=pd_threatened,col="red",lwd=3) # v= 179194.2; vertical line in red showing actual loss
abline(v=mean(total_pd-pd),col="blue",lwd=3)# mean random pd loss = 139083.2; vertical line in blue showing the mean of random loss
CI <- quantile(total_pd-pd,prob=c(0.025,0.975)) # confidence interval
#CI = [136766.0; 141371.8] 

######
```
