## Supplementary figures and images for "Over half of the phylogenetic diversity accumulated on the African plant tree of life may be eroded under current biodiversity crisis even though threatened species are not evolutionarily unique"

### Figure S1.pdf

#### Map showing sampling efforts

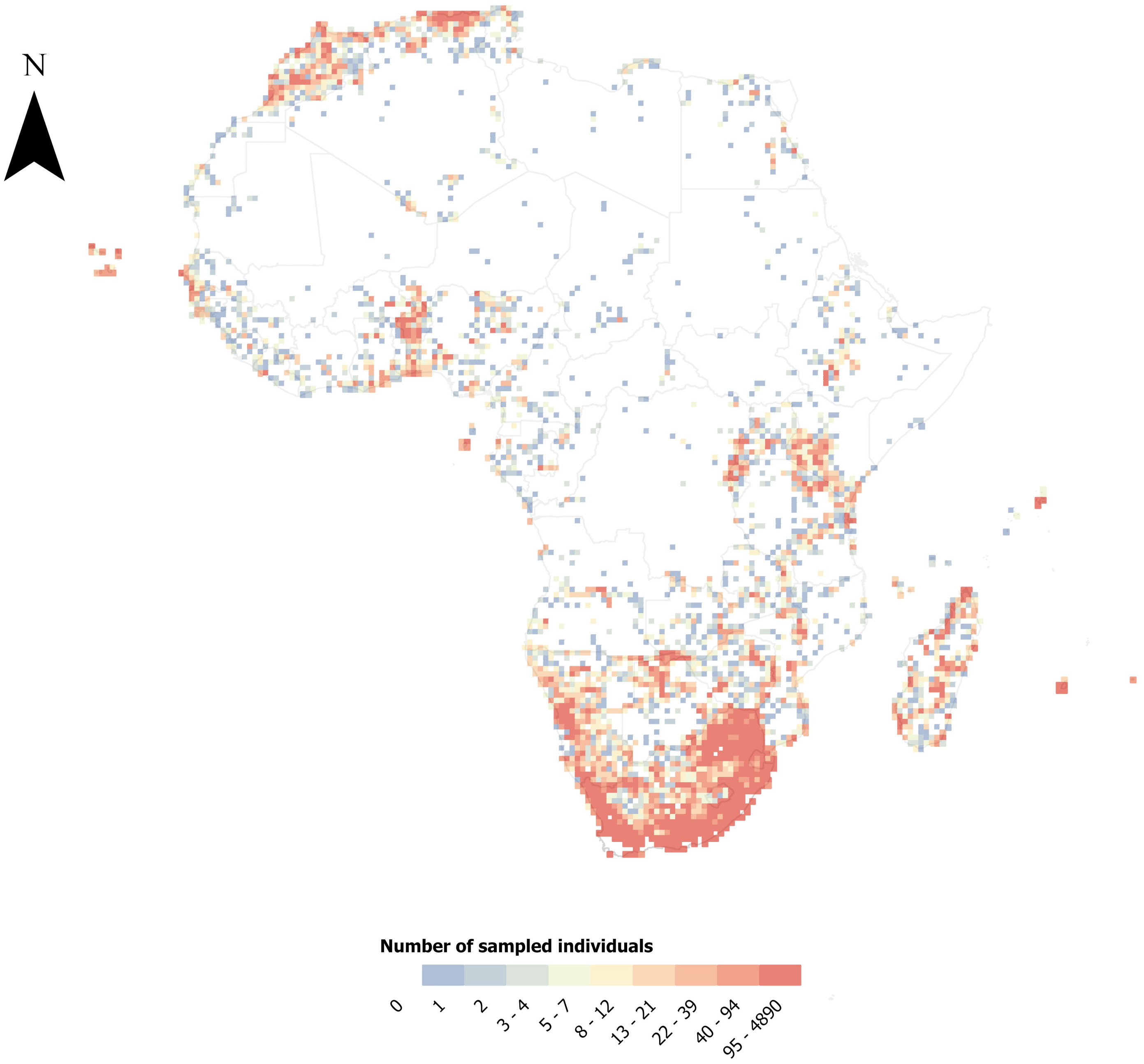

### Figure S2.pdf

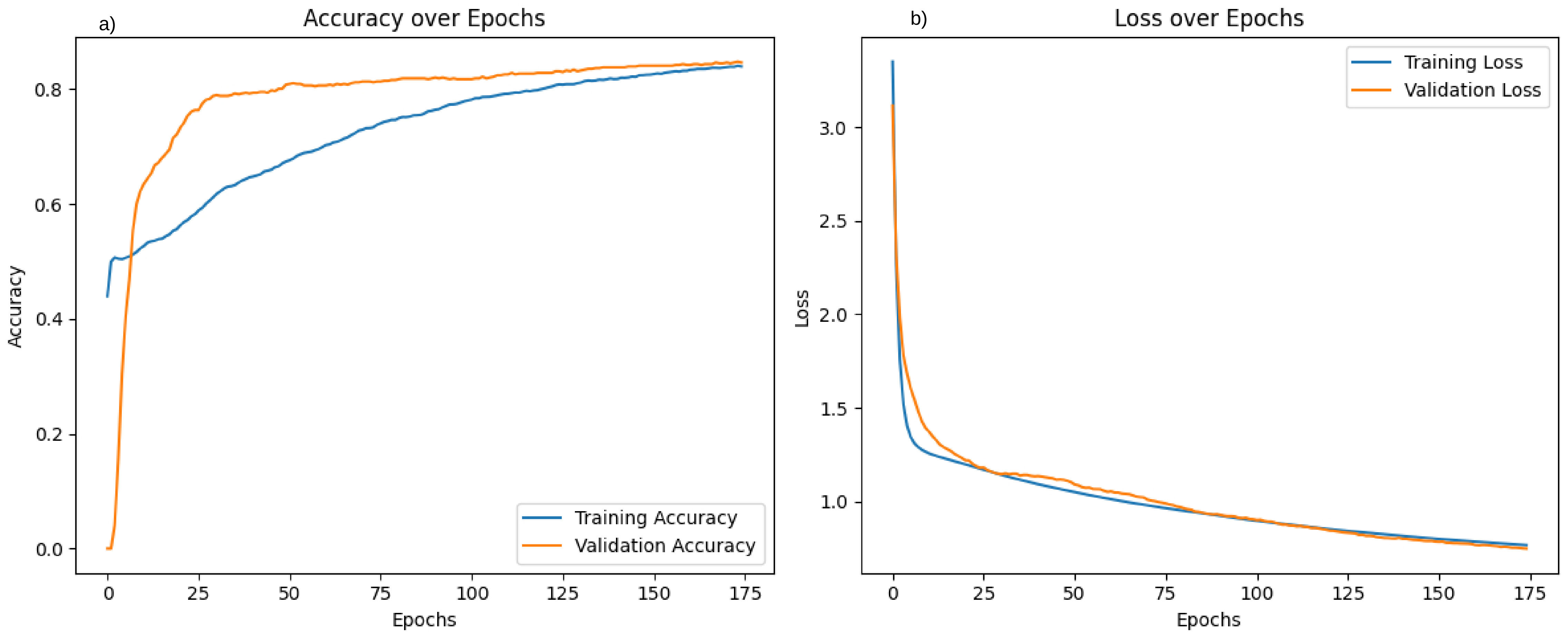

### Figure S3.pdf

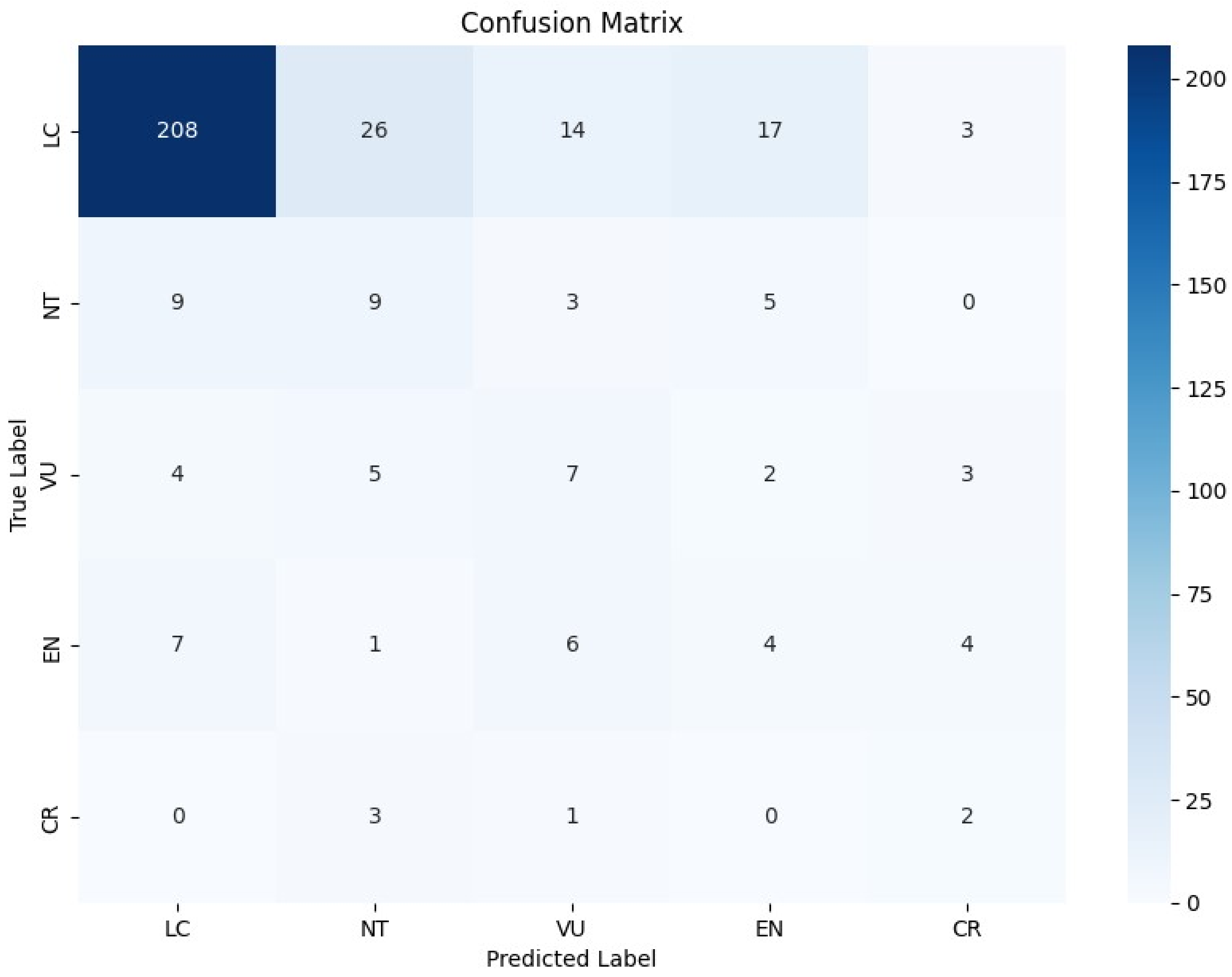

### Figure S5.pdf

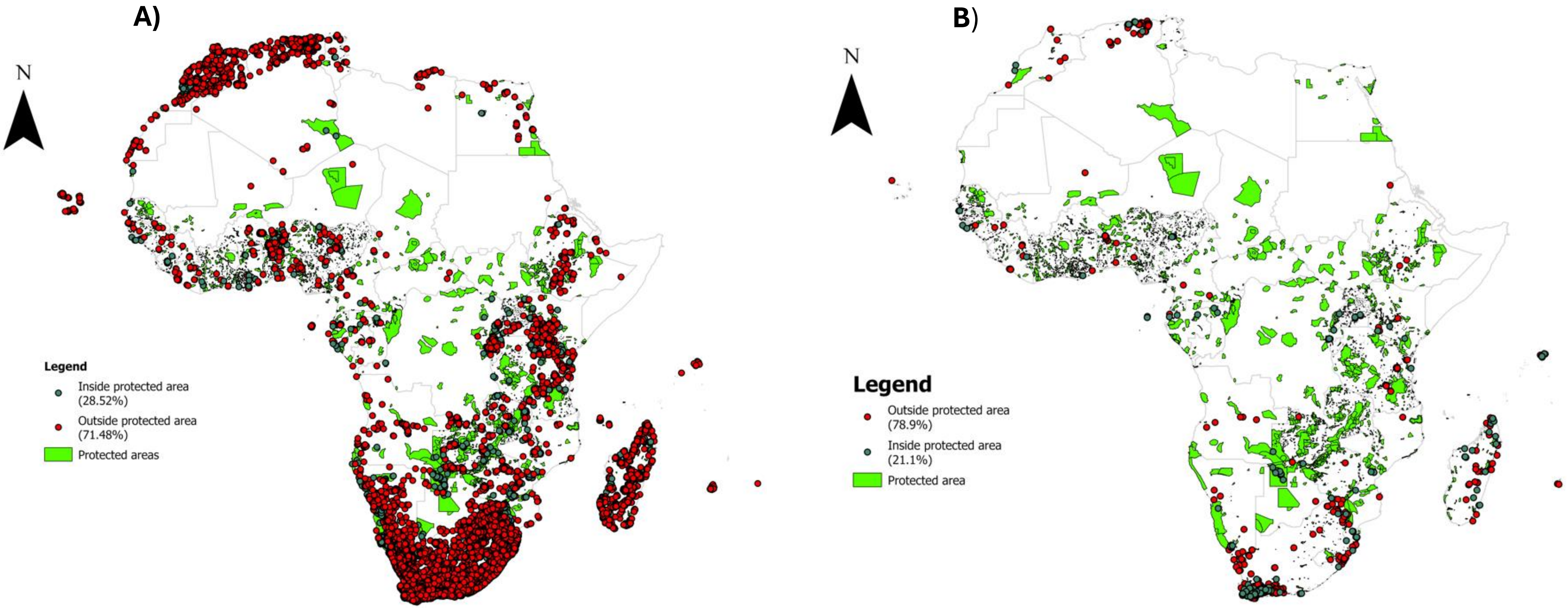
